# Tricyclic and tetracyclic antidepressants upregulate VMAT2 activity and rescue disease-causing VMAT2 variants

**DOI:** 10.1101/2023.10.09.561601

**Authors:** Xunan Wang, Ilias Marmouzi, Peter SB Finnie, Svein I Støve, Meghan L Bucher, Tatiana V Lipina, Amy J Ramsey, Gary W Miller, Ali Salahpour

## Abstract

Vesicular monoamine transporter 2 (VMAT2) is an essential transporter that regulates brain monoamine transmission and is important for mood, cognition, motor activity, and stress regulation. However, VMAT2 remains underexplored as a pharmacological target. In this study, we report that tricyclic and tetracyclic antidepressants acutely inhibit, but persistently upregulate VMAT2 activity by promoting VMAT2 protein maturation. Importantly, the VMAT2 upregulation effect was greater in BE(2)-M17 cells that endogenously express VMAT2 as compared to a heterologous expression system (HEK293). The net sustained effect of tricyclics and tetracyclics is an upregulation of VMAT2 activity, despite their acute inhibitory effect. Furthermore, imipramine and mianserin, two representative compounds, also demonstrated rescue of nine VMAT2 variants that cause Brain Vesicular Monoamine Transport Disease (BVMTD). VMAT2 upregulation could be beneficial for disorders associated with reduced monoamine transmission, including mood disorders and BVMTD, a rare but often fatal condition caused by a lack of functional VMAT2. Our findings provide the first evidence that small molecules can upregulate VMAT2 and have potential therapeutic benefit for various neuropsychiatric conditions.

## Introduction

Monoamine regulation in the central nervous system (CNS) is important for cognition, motor control, mood, arousal, stress, and temperature regulation [1–3]. Altered monoamine transmission is hypothesized to be one of the underlying causes of depression and anxiety [4, 5]. Vesicular monoamine transporter 2 (VMAT2) is an important transporter for monoamine transmission within the CNS and plays a central role in synaptic vesicular packaging of monoamine neurotransmitters, which include serotonin, dopamine, norepinephrine, epinephrine, and histamine [6–7]. Rare homozygous variants in the *SLC18A2* gene, which encodes the VMAT2 protein, result in brain vesicular monoamine transport disease (BVMTD) [8–9]. Symptoms range from global developmental delay and hypotonia in mild cases to childhood lethality in severe cases [8–9]. In addition, lower VMAT2 levels are associated with the risk of Parkinson’s disease and VMAT2 inhibition by reserpine induces depression in humans [10–15].

Since the 1950s, the monoamine system has been targeted pharmacologically for depression and mood disorders by monoamine oxidase inhibitors (MAOIs) and tricyclic antidepressants (TCAs) and since the 1980s by selective serotonin reuptake inhibitors (SSRIs) [16]. However, unlike plasma membrane transporters for serotonin (SERT) and norepinephrine (NET), the established targets for TCAs, VMAT2 has remained an underexplored pharmacological target for mood disorders and other indications [7, 16].

Our study aimed to identify compounds that could upregulate VMAT2 levels and activity since low VMAT2 levels are associated with BVMTD, Parkinson’s disease, and depression, and high VMAT2 levels are protective for Parkinson’s disease risk in humans and neurotoxicity in rodents [8–15, 17–18]. We hypothesized that tricyclic antidepressants modulate VMAT2 activity since they are known to affect the monoamine transporters SERT and NET. We also tested the effects of tetracyclic antidepressants on VMAT2 activity because they are structurally similar to tricyclics and are used for the same clinical indications. Our results show that acute treatments with tricyclic and tetracyclic antidepressants inhibit VMAT2 activity, while sustained treatments upregulate VMAT2 protein levels and activity. Furthermore, our results show that these compounds can rescue the activity of nine BVMTD-causing missense VMAT2 variants that impair VMAT2 function. Overall, our study is the first to report upregulation of VMAT2 protein and activity using clinically-effective tricyclic and tetracyclic antidepressants.

## Materials and Methods

The suppliers of reagents are listed in **Table S1**.

### Plasmids

The full-length mouse and human *SLC18A2* cDNAs encoding VMAT2 were cloned into pcDNA3.1 vectors. A YFP-HA-DAT plasmid encoding WT human dopamine transporter (DAT) with a C-terminal EYFP tag and an HA-tag in the second extracellular loop (in pEYFP-C1 vector) was a gift from Sorkin Lab, University of Pittsburgh [19]. Constructs for human VMAT2 BVMTD variants were cloned by PCR-based site-directed mutagenesis using primer pairs in **Table S2** [20].

### Cell lines

Cell lines were maintained in a 37℃ incubator with 5% CO_2_. HEK293, HEK293T, and BE(2)-M17 cells were obtained from ATCC. The HEK-VMAT2 cell line was generated by stably transfecting the mouse VMAT2 cDNA expression construct in HEK293 cells. The HEK-mCherry-VMAT2 cells were generated previously [21]. The HEK-YFP-DAT stable cell line was generated previously [22]. All cell lines were maintained in DMEM medium supplemented with 10% fetal bovine serum, 100units/mL penicillin, and 100μg/mL streptomycin. Stable cell lines were additionally supplemented with 0.5mg/ml G418 Sulfate.

### FFN206 uptake assay

Black-bottomed 96-well plates were precoated overnight with 0.1mg/ml poly-D-lysine and UV sterilized before use. All incubations and washes were done at 37℃.

For the inhibition assay, HEK-VMAT2 cells were seeded at 50,000 cells/well on day 1. On day 2, the growth medium was aspirated and 90μL of drug-containing medium (1% DMSO, no G418) at varying concentrations were added to each well and incubated for 30min. After the incubation, 10μL of false fluorescent neurotransmitter (50μM FFN206, diluted in PBS) was added to each well, resulting in a 5μM final concentration of FFN206. Cells were incubated with FFN206 for 1hr. FFN206 uptake was terminated by aspirating the medium, and cells were washed twice with 200μL warm PBS (supplemented with MgCl_2_ and CaCl_2_), each with 5min incubation. Subsequently, 100μL warm PBS was added to each well and the plate was read with a Mithras LB 940 plate reader at excitation/emission filters 380nm/450nm.

For the upregulation assay, HEK-VMAT2 cells were seeded at 30,000 cells/well on day 1. On day 2, the growth medium was aspirated and 100uL of medium containing drug or vehicle (1% DMSO, no G418) was added to each well and incubated for 18hrs. On day 3, the drug-containing medium was aspirated and washed 3 times with 200μL warm medium, each with 20min incubation. After the wash, 90μL medium containing 100μM tetrabenazine (TBZ) and 1% DMSO or vehicle were added to each well and incubated for 30min. After incubation, 10μL of 50μM FFN206 diluted in PBS was added to each well and incubated for 1hr. Afterwards the FFN206 uptake was terminated by 2 washes with 200μL warm PBS, each with 5min incubation. Subsequently, 100μL warm PBS was added to each well and the plate was read with a Mithras LB 940 plate reader. ***Epifluorescent Microscopy***

FFN206 uptake was performed as previously described [21]. Briefly, HEK-mCherry-VMAT2 cells were seeded in a clear 6-well plate at 250,000 cells/well density. 24hrs later, culture media was removed and cells were pre-incubated with DMSO or 10μM TBZ for 30min in DMEM. After 30min, FFN206 prepared in DMEM was added to a final concentration of 5μM and incubated for 1hr. Afterwards, media was removed and replaced with DMEM containing 1:25 trypan blue to reduce background fluorescence and cells were imaged on an Olympus APEX100 microscope at 40x magnification using a DAPI filter to visualize FFN206 and mCherry/TexasRed filter to visualize mCherry-tagged VMAT2.

### Flow cytometry

BE(2)-M17 cells were seeded at 100,000 cells/well density in clear 24-well plates. On day 2, the growth medium was aspirated and 500μL of 10μM drug-containing medium (0.1% DMSO) or vehicle were added to each well and incubated for 18hrs. On day 3, the drug-containing medium was aspirated and washed 3 times with 500μL warm medium, each with 20min incubation. After the wash, 450μL medium containing 100μM reserpine and 1% DMSO or vehicle were added to each well and incubated for 30min. After incubation, 50μL of 50μM FFN206 diluted in PBS was added to each well and incubated for 1hr. The FFN206 uptake was terminated by a 1ml warm PBS wash and then trypsinized with 200μL trypsin+EDTA (0.25%). The cells were then resuspended in 400μL flow cytometry buffer (25mM HEPES, pH7.0, 1% bovine serum albumin, 1mM EDTA, in 1X PBS) and kept on ice. Flow cytometry was performed under operator assistance at the University of Toronto Flow Cytometry Facility using BD LSR Fortessa™ X-20 flow cytometer with a 355nm laser and a 450/50nm filter. The flow cytometry data was analyzed with the FlowJo software. Example gating can be found in **Figure S1**. The mean fluorescence intensity (MFI) of the FFN+ population was quantified and normalized to cells that were treated with vehicle.

### Western blot

HEK293, HEK-VMAT2, and HEK-YFP-DAT cells were seeded at 5,000,000 cells/well density in 10cm tissue culture dishes. The next day, the growth medium was aspirated and 10mL of medium (0.1% DMSO) containing 50μM and 100μM imipramine or mianserin or vehicle were added to the cells and incubated for 18hrs. On the collection day, cells were washed once with cold PBS, then harvested with 2mL Tris-EDTA buffer (25mM Tris-HCl, 1mM EDTA, pH7.4) containing freshly-added protease/phosphatase inhibitors. The cells were mechanically lysed on ice by Polytron (medium setting, 5sec on, 5sec off, two cycles), and centrifuged at 800xg for 10min at 4℃ to pellet the nuclear fraction. The resulting supernatant was transferred into a new set of tubes and centrifuged again at 29,097xg for 40min at 4℃. The supernatant was discarded and the pellet containing membrane proteins was resuspended with Tris-EDTA buffer with inhibitors and 0.2% SDS by vortexing until the pellet dissolves. The protein concentration was determined using Pierce™ BCA Protein Assay Kits following the manufacturer’s instructions.

For the deglycosylation experiment, 22μg of vehicle-treated VMAT2 protein lysate were digested with EndoH, PNGase F, or reaction buffer following manufacturer’s non-denaturing condition protocol for 24hrs.

Samples were heated at 55℃ for 10min before proteins were loaded in each well. SDS-PAGE was performed using NuPAGE™ 4-12% Bis-Tris Mini Protein Gel and MOPS-SDS running buffer (50mM MOPS, 50mM Tris, 0.1% SDS, 1mM EDTA). The proteins were transferred onto a 0.45nm PVDF membrane, and the total protein was stained using Revert™ 700 Total Protein Stain. The total protein was visualized at 700nm on an Odyssey® M imaging system (LI-COR, Inc.). The membrane was blocked with 3% non-fat milk in TBST (19mM Tris, 137mM NaCl, 2.7mM KCl, 0.1% Tween 20) for 1hr and then incubated in primary antibodies diluted in 3% non-fat milk in TBST. For the VMAT2 blot, the membrane was incubated with rabbit serum (in 50% glycerol) containing anti-mVMAT2 antibody against C-terminal peptide (TQNNVQPYPVGDDEESESD) at 1:3000 dilution overnight at 4℃ [23]. For the YFP-DAT blot, the membrane was incubated with rabbit-anti-GFP antibody (A11122) at 1:1500 dilution for 2hrs at room temperature. IRDye® 800CW Goat anti-Rabbit IgG secondary antibody at 1:7000 dilution in 3% non-fat milk in TBST was used for both VMAT2 and YFP-DAT blots. The blots were visualized under 700nm and 800nm channels on the Odyssey® M imaging system (LI-COR, Inc.) and quantified using ImageJ.

### Transient transfection of BVMTD variants

On day 1, HEK293T cells were seeded at 50,000 cells/well density in PDL-precoated black-bottomed 96-well plates. 4 hours after seeding, 50ng of human VMAT2 (WT or BVMTD variant) plasmid and 50ng of pcDNA3.1 empty plasmid were co-transfected into each well using Lipofectamine^TM^ 3000, following the manufacturer’s protocol for 96-well plate experiments. 22 hours post transfection, old media was aspirated and either 50μM of drug (imipramine or mianserin)-containing media (0.05% DMSO) or vehicle (0.05% DMSO) was added to the cells. 18hrs after, the FFN206 uptake assay was performed as described in the previous section for upregulation assay.

### RT-qPCR

HEK-VMAT2, HEK-YFP-DAT, and BE(2)-M17 cells were seeded in 6-well tissue culture plates at 1,000,000 cells/well density. The next day, the growth medium was aspirated and 10mL of medium (0.1% DMSO) containing 100μM (for HEK-VMAT2 and HEK-YFP-DAT cells) or 10μM (for BE(2)-M17 cells) imipramine or mianserin or vehicle were added to the cells and incubated for 18hrs. On the third day, media was aspirated and the cells were directly collected in 1ml/well of Trizol for RNA isolation [24]. RNA was reverse-transcribed into cDNA using the SuperScript™ IV VILO™ Master Mix with ezDNase™ Enzyme. Quantitative RT-qPCR was performed using PowerUp™ SYBR™ Green Master Mix and QuantStudio 3 Real-Time PCR System (ThermoFisher Scientific).

Primer pairs used for quantitative PCR are listed in **Table S3**. The relative fold of gene expression was calculated as 2^-(ΔΔCt) normalized to GAPDH levels.

## Results

### Acute treatment with tricyclics or tetracyclics inhibits VMAT2 activity

FFN206 is a fluorescent substrate of VMAT2 that gets transported into VMAT2-containing intracellular compartments such as vesicles [21, 25]. We detected colocalization of VMAT2 and FFN206 using epifluorescent microscopy (**Figure S2**) and used FFN206 uptake to measure VMAT2 activity in HEK cells stably expressing mouse VMAT2 (HEK-VMAT2). We tested seven tricyclics and three tetracyclics that are approved for clinical use in humans (**Figures 1A-B**). The acute effect of the drugs on VMAT2 activity was first assessed by pre-incubating HEK-VMAT2 cells with test compounds for thirty minutes followed by FFN206 co-incubation. Thirty-minute treatment with 10μM of all tested tricyclics or tetracyclics significantly inhibited VMAT2 activity, except for the tetracyclic mirtazapine (**Figures 1C-D**). Next, we assessed the potency of the tricyclics and tetracyclics using dose response experiments. All drugs demonstrated comparable dose-dependent inhibition of VMAT2 with low-to-mid micromolar potency (IC_50_), except for mirtazapine (**Figures 1E-F**). Protriptyline (IC_50_ = 3.88μM) and amoxapine (IC_50_ = 5.29μM) were the most potent tricyclic and tetracyclic, respectively (**Table 1**). Compared to tetrabenazine (TBZ), the prototypical VMAT2 inhibitor, tricyclics and tetracyclics inhibited VMAT2 with a lower potency (higher IC_50_) but similar maximum inhibition (I_max_) (**Table 1**).

**Figure 1.**
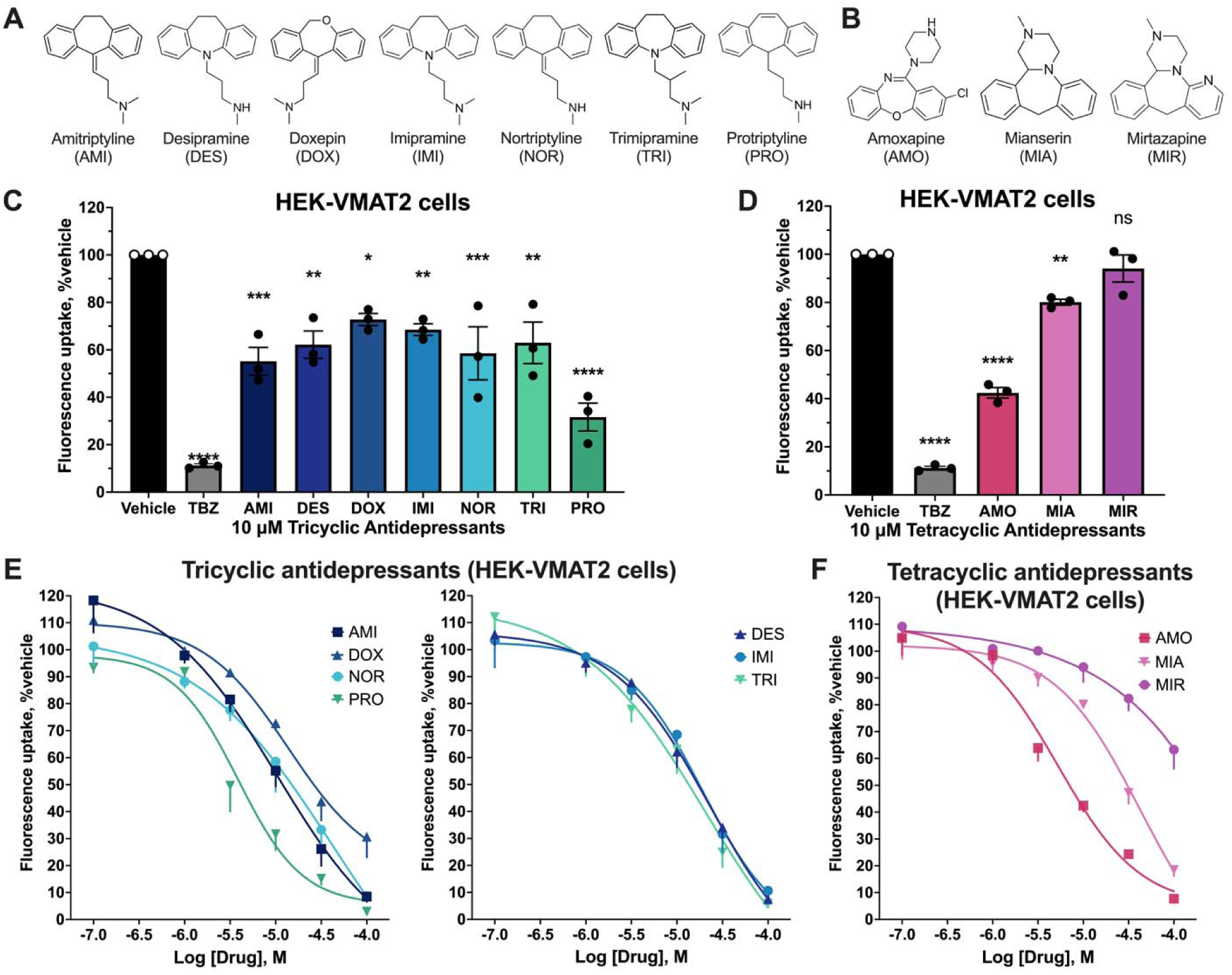
Acute effect of tricyclics and tetracyclics on VMAT2 activity. **(A)** Chemical structures of tricyclics tested. **(B)** Chemical structures of tetracyclics tested. **(C, D)** Effect of 30-minute incubation with 10μM tricyclics or tetracyclics on VMAT2 activity in HEK-VMAT2 cells. One-way ANOVA was performed followed by Dunnett’s test comparing each treatment against the vehicle (n = 3). P > 0.05 (ns), ≤ 0.05 (*), ≤ 0.01 (**), ≤ 0.001 (***), ≤ 0.0001 (****). **(E, F)** Dose-response of 30-minute incubation with tricyclics and tetracyclics on VMAT2 activity in HEK-VMAT2 cells (n = 3). For each independent experiment, all treatments were normalized to vehicle (100%).

**Table 1.** Comparison of VMAT2 inhibition and upregulation effects by tricyclics and tetracyclics.

|  |  | Inhibition (30min) |  | Upregulation (18hrs) |  |  | IC<sub>50</sub>/EC<sub>50</sub> |
| --- | --- | --- | --- | --- | --- | --- | --- |
|  |  | IC<sub>50</sub> (μM) | I<sub>max</sub>, %<br>(mean ± SEM) | EC<sub>50</sub> (μM) | E<sub>max</sub>, %<br>(mean ± SEM) | E<sub>max</sub><br>concentration<br>(μM) |  |
| **Tricyclics** | **AMI** | 10.46 | 8.47±1.88 | 314.90* | 237.26±7.63 | 31.62 | * |
|  | **DES** | 22.63 | 7.51±1.28 | 17.29 | 177.59±5.96 | 56.23 | 1.31 |
|  | **DOX** | 13.78 | 30.72±7.66 | 41.91 | 243.49±19.69 | 100 | 0.33 |
|  | **IMI** | 19.05 | 10.67±2.24 | 30.66 | 245.36±15.06 | 100 | 0.62 |
|  | **NOR** | 40.69 | 8.73±1.11 | 8884* | 207.90±18.90 | 31.62 | * |
|  | **TRI** | 20.66 | 5.87±1.33 | 31.06 | 194.63±22.32 | 100 | 0.67 |
|  | **PRO** | 3.88 | 2.93±0.24 | 6845* | 223.92±11.02 | 31.62 | * |
| **Tetracyclics** | **AMO** | 5.29 | 7.73±1.23 | 17.88 | 179.83±18.89 | 31.62 | 0.30 |
|  | **MIA** | 42.32 | 18.41±2.14 | 29.50 | 223.48±8.24 | 100 | 1.44 |
|  | **MIR** | 4629000* | 63.27±7.1 | 33.68 | 175.11±7.95 | 100 | * |
| **Control** | **TBZ** | 0.01** | 9.51±0.67 |  |  |  |  |

\*Cannot be estimated accurately because of incomplete logarithmic dose-response curves.
\*\*Estimated using the complete dose-response curve in **Figure S3**.

### Sustained treatment with tricyclics or tetracyclics upregulates VMAT2 activity

With the knowledge that some inhibitors could also act as pharmacological chaperones of the given target and increase its protein maturation and levels, we then tested the sustained effect of each tricyclic and tetracyclic on VMAT2 activity after an 18-hour treatment [22]. To prevent the presence of inhibitors masking a VMAT2 upregulation effect (measured by FFN206 uptake activity), we incorporated three twenty-minute washes to remove any remaining drug after the treatment. In HEK-VMAT2 cells, 18-hour treatment with the majority of evaluated tricyclics, including desipramine, imipramine, nortriptyline, trimipramine, and protriptyline, caused significant upregulation of VMAT2 activity at 10μM concentration (**Figure 2A**). In contrast, at 10μM concentration, no sustained tetracyclics treatment significantly upregulated VMAT2 activity (**Figure 2B**). We next conducted dose response curves in which all tricyclics and tetracyclics demonstrated a dose-dependent upregulation of VMAT2 with mid-micromolar to low-millimolar potency (EC_50_) (**Figures 2C-D**). Desipramine (EC_50_ = 17.29μM) and amoxapine (EC_50_ = 17.88μM) were the most potent upregulators among tricyclics and tetracyclics, respectively (**Table 1**). Sustained treatment with 100μM imipramine and mianserin resulted in the highest VMAT2 activity upregulation of their class at an E_max_ of 245% and 233%, respectively (**Table 1**). To assess whether VMAT2 upregulation is only seen in an over-expression system, we carried out sustained treatment experiments on BE(2)-M17 cells, which endogenously express low levels of human VMAT2. We used flow cytometry to measure uptake of FFN206 by BE(2)-M17 cells because the endogenous levels of VMAT2 were below the detection limit of the plate reader used with HEK cells. Reproducible and larger-fold VMAT2 upregulation was observed in BE(2)-M17 cells after sustained treatment with tricyclics and tetracyclics. At 10μM concentrations, all tricyclics and tetracyclics demonstrated significant upregulation of VMAT2 activity ranging from 152% to 498% of vehicle (**Figures 2E-F**).

**Figure 2.**
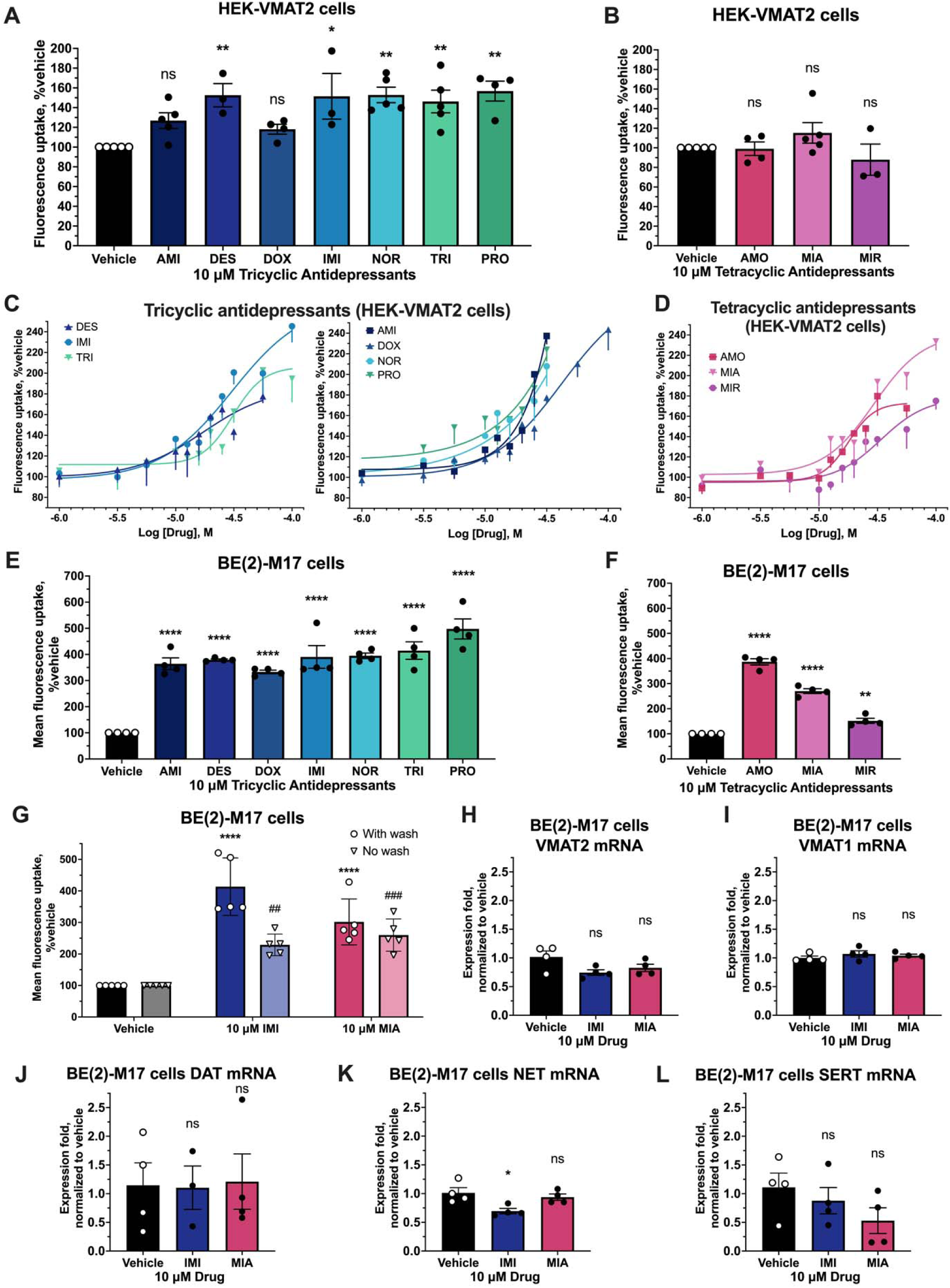
Sustained effect of tricyclics and tetracyclics on VMAT2 activity. **(A, B)** Effect on VMAT2 activity of 18-hour incubation with 10μM tricyclics or tetracyclics followed by washout in HEK-VMAT2 cells. One-way ANOVA was performed followed by Dunnett’s test comparing each treatment against the vehicle (n = 3-6). **(C, D)** Dose-response effect on VMAT2 activity of 18-hour incubation with each tricyclic or tetracyclic at 10μM, followed by washout in HEK-VMAT2 cells (n = 3-6). **(E, F)** Effect on endogenous VMAT2 activity of 18-hour incubation with each tricyclic or tetracyclic at 10μM, followed by washout in BE(2)-M17 cells (n = 4). One-way ANOVA was performed followed by Dunnett’s test comparing each treatment against the vehicle. (**G**) Effect on endogenous VMAT2 activity of 18-hour incubation with 10μM imipramine and mianserin with or without washout in BE(2)-M17 cells (n = 5). (**H**-**L**) RT-qPCR of endogenous monoamine transporters mRNA in BE(2)-M17 cells after 18-hour treatment with 10μM imipramine or mianserin (n = 4). Two-way ANOVA was performed followed by Dunnett’s test comparing between drug and vehicle for each wash condition. P > 0.05 (ns), ≤ 0.05 (* or ^#^), ≤ 0.01 (** or ^##^), ≤ 0.001 (*** or ^###^), ≤ 0.0001 (**** or ^####^). For each independent experiment, all treatments were normalized to vehicle (100%).

To delve into the upregulation effect of tricyclics and tetracyclics, we selected a representative drug candidate for each antidepressant class based on the dose-response of inhibition and upregulation in HEK-VMAT2 cells. An IC_50_/EC_50_ ratio was calculated to estimate the relative inhibition-to-upregulation potency (**Table 1**). IC_50_/EC_50_ <1 indicates that the compound is more potent as an inhibitor and IC_50_/EC_50_ > 1 indicates that the compound is more potent as an upregulator. Imipramine was chosen as the representative tricyclic because it had the highest E_max_ within the class and a relatively large IC_50_/EC_50_ ratio. Mianserin was chosen as the representative tetracyclic because it had both the highest E_max_ for upregulation within the class and the largest IC_50_/EC_50_ ratio. Because a larger upregulation effect of tricyclics and tetracyclics was seen in BE(2)-M17 cells compared to HEK-VMAT2 cells, we hypothesized that, under sustained treatment, the upregulation effect would outweigh the inhibitory effect. We therefore tested the net effect of sustained imipramine or mianserin treatment with and without drug washout. As shown in **Figure 2G**, sustained treatment without washout still lead to a significant upregulation of endogenous VMAT2 activity in BE(2)-M17 cells.

We next investigated whether the upregulation of VMAT2 activity by tricyclics or tetracyclics could be explained by upregulation of gene expression of VMAT2 or related monomaine transporters (VMAT1, DAT, NET, and SERT) in BE(2)-M17 cells, which express these transcripts endogenously. With 10μM imipramine or mianserin treatment, despite a significant upregulation in activity was observed, no upregulation in any monoamine transporter mRNA, including VMAT2 was detected (**Figures 2E-F, H-L**). ***Tricyclics and tetracyclics upregulate VMAT2 by promoting VMAT2 protein maturation*.**

We and others have observed that some transporter ligands can act as pharmacological chaperones that increase the ratio of mature:immature transporter [22, 26–29]. Therefore, we investigated whether drug treatment affected the process of VMAT2 protein maturation as a potential mechanism explaining the upregulation of VMAT2 activity by tricyclics and tetracyclics. As expected for a glycosylated transmembrane protein, antibodies against VMAT2 revealed proteins with molecular weights of 41, 50, 60, and 80kDa in HEK-VMAT2 cells (**Figure 3A**). We treated cells with deglycolysating enzymes to differentiate the immature and fully glycosylated isoforms of VMAT2. Deglycosylation by endoglycosidase H (Endo H) revealed that the 50kDa band, which was sensitive to Endo H digestion, represents the immature VMAT2 located at the endoplasmic reticulum (ER) [22] (**Figure 3A**). Deglycosylation by peptide:N-glycosidase F (PNGase F) revealed that both 60kDa and 80kDa bands, which were insensitive to Endo H digestion but sensitive to PNGase F digestion, represent the mature VMAT2 protein (**Figure 3A**).

**Figure 3.**
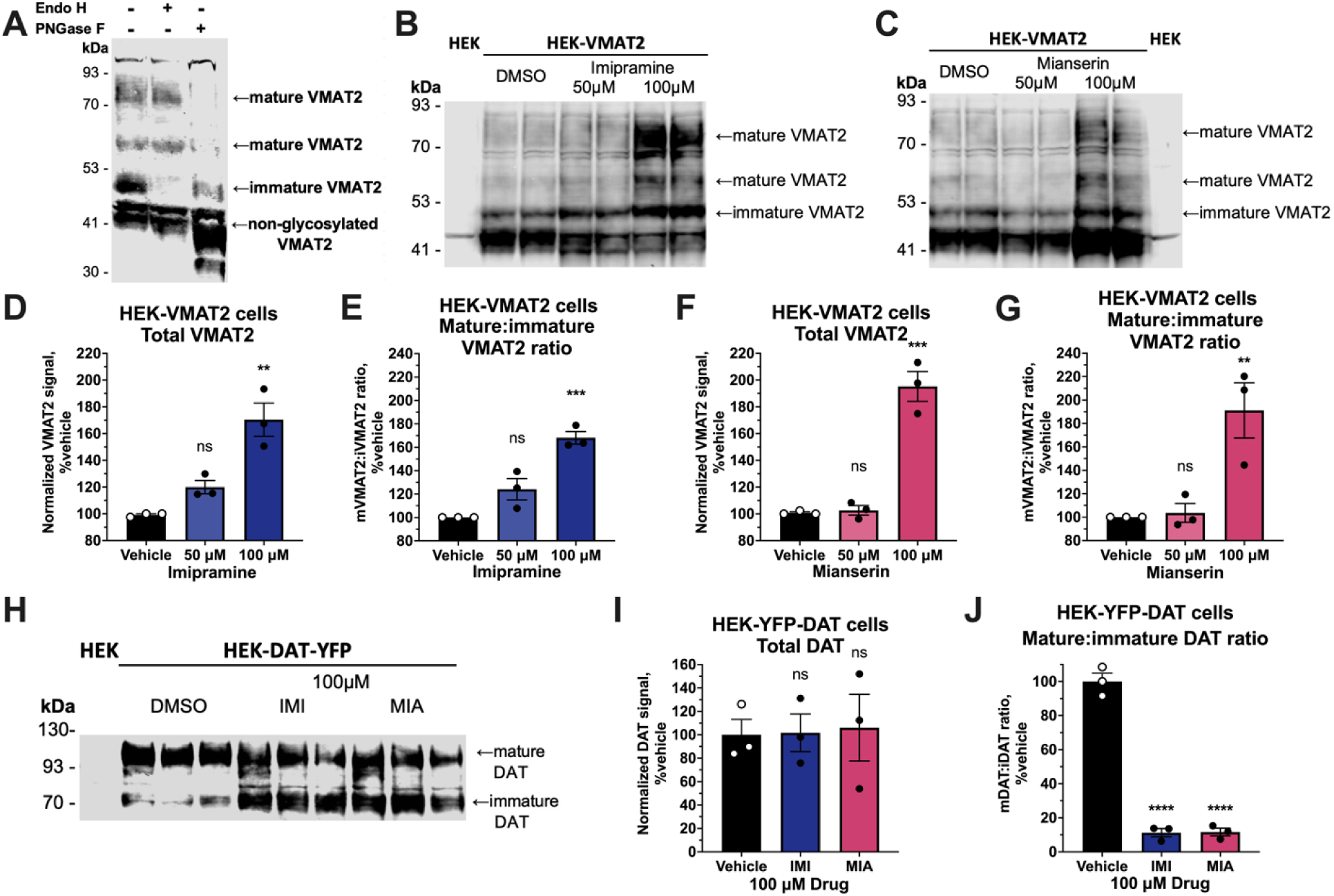
Sustained effect of tricyclics and tetracyclics on VMAT2 protein and mRNA. **(A)** Western blot of endoglycosidase H (EndoH) and peptide:N-glycosidase F (PNGase F) digested VMAT2 protein revealing different trafficking stages of the VMAT2 protein. (**B, C**) Representative western blot of 18-hour imipramine or mianserin incubation on VMAT2 protein in HEK-VMAT2 cells. Duplicate wells (n=1) displayed. (**D-G**) Quantification of western blot in (**B, C**) (n=3). (**H**) Western blot for DAT protein after 18-hour 100 μM imipramine or mianserin incubation in HEK-YFP-DAT cells (n = 3, each well represents an independent experiment). (**I, J**) Quantification of western blot in (**I**) (n = 3). For all bar graphs, one-way ANOVA was performed followed by Dunnett’s test comparing each treatment against the vehicle. P > 0.05 (ns), ≤ 0.05 (*), ≤ 0.01 (**), ≤ 0.001 (***), ≤ 0.0001 (****).

We next treated the HEK-VMAT2 cell line with imipramine and mianserin and determined the ratio between mature VMAT2 and immature VMAT2, where a higher ratio represents a greater extent of VMAT2 maturation. Western blot analysis of cells treated for 18-hours with 100μM imipramine or mianserin both showed a significant increase in total VMAT2 protein and mature:immature VMAT2 ratio (**Figures 3B-G**). The increase in mature:immature VMAT2 ratio suggests that imipramine and mianserin promote VMAT2 maturation. We were unable to assess the effects of compounds on VMAT2 protein levels in BE(2)-M17 cells because the VMAT2 protein levels in these cells were below the western blot detection threshold (data not shown).

To examine the selectivity of these compounds, we assessed the effect of imipramine and mianserin on DAT (*SLC6A3*) by western blot of stably transfected HEK-293 cells. Like VMAT2, functional, mature DAT protein is glycosylated and is detected by western blot in its mature (110kDa) and immature (70kDa) state (**Figures 3H**) [22]. In contrast to the effects on VMAT2 protein, 18-hours treatment with 100μM imipramine nor mianserin did not affect total DAT protein and significantly decreased the mature:immature DAT ratio (**Figures 3H-J**).

### Imipramine and mianserin rescue the activity of missense VMAT2 BVMTD variants

To explore potential clinical applications of VMAT2 upregulation by tricyclics and tetracyclics, we tested their effects on nine disease-causing, missense VMAT2 variants that were recently reported [8, 30]. All nine missense variant showed impaired VMAT2 uptake activity when studied in transiently-transfected HEK293T cells (**Figure 4A**). We then tested the effect of 18-hour treatment with 50μM imipramine or mianserin on all variants. Both imipramine and mianserin caused a significant upregulation of activity of all VMAT2 variants (**Figures 4B-K**). The upregulation effect ranged from 179% to 321% of vehicle treatment. For most variants, the effect on VMAT2 activity was greater with imipramine than mianserin.

**Figure 4.**
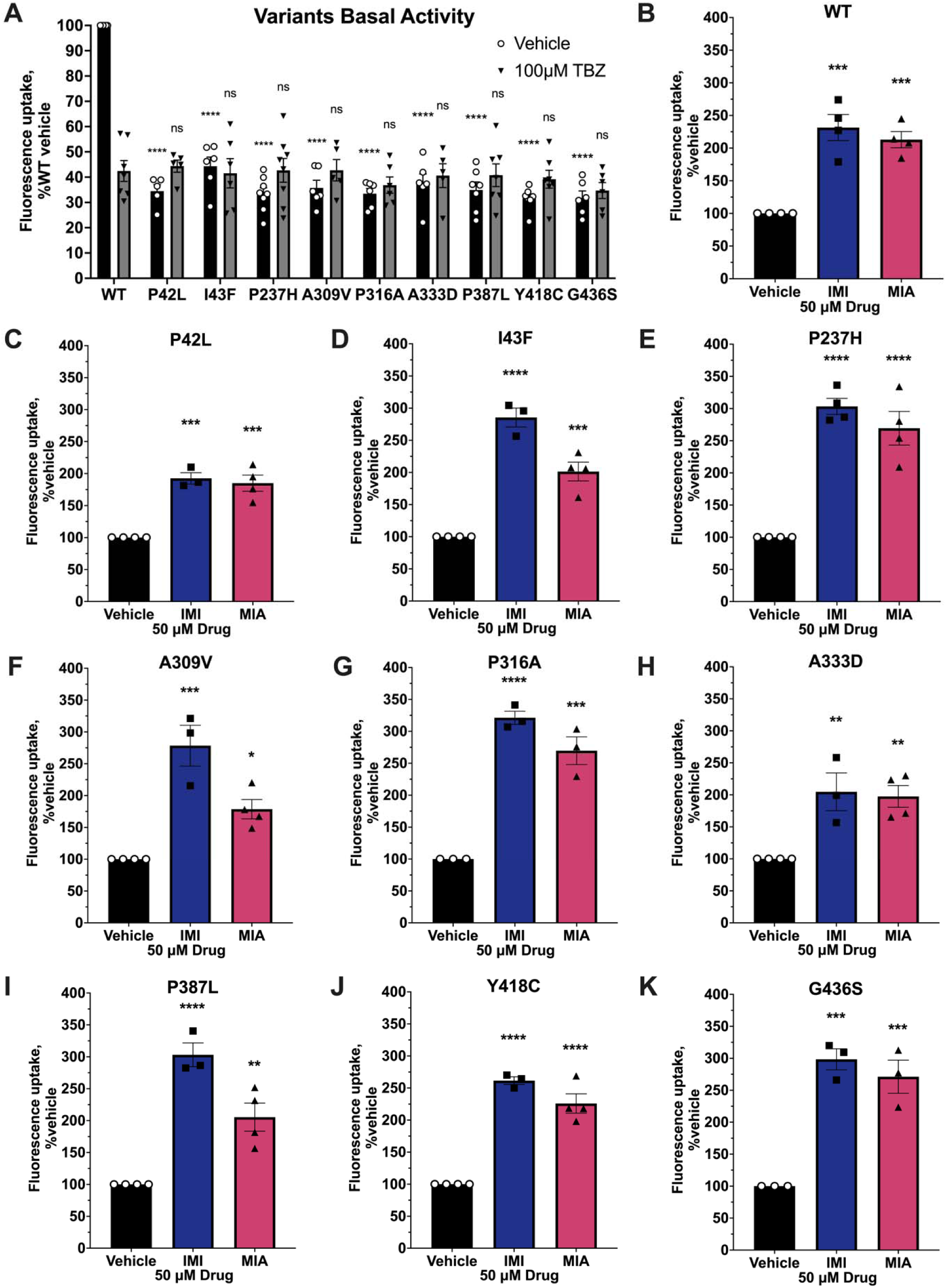
Sustained effect of imipramine and mianserin on the activity of VMAT2 BVMTD variants. (**A**). Basal VMAT2 activity of transiently transfected BVMTD variants in HEK293T cells (n = 5-8). Two-way ANOVA was performed followed by Dunnett’s test comparing each variant against WT for both vehicle and 100μM TBZ conditions. (**B-K**) Transiently transfected VMAT2 WT and variant activity after 18-hour treatment with 50 μM or 100 μM imipramine and mianserin, respectively (n = 3-4). One-way ANOVA was performed followed by Dunnett’s test comparing each treatment against the vehicle. P > 0.05 (ns), ≤ 0.05 (*), ≤ 0.01 (**), ≤ 0.001 (***), ≤ 0.0001 (****). For each independent experiment, all treatments were normalized to vehicle (100%).

## Discussion

We report that VMAT2 is a novel pharmacological target of several clinically-relevant tricyclic and tetracyclic antidepressants. With an apparent class effect, these antidepressants exhibit two modes of VMAT2 modulation: inhibition of VMAT2 activity with acute treatment, and upregulation of VMAT2 activity with sustained treatment. The mechanism of inhibition could be explained by a direct interaction between the drug and the target, causing inhibition of VMAT2 transporter activity. The mechanism of upregulation includes at least in part an effect on protein level and maturation. In BE(2)-M17 cells that endogenously express VMAT2 and related monoamine transporter transcripts, a drug concentration that result in 390% and 270% upregulation of VMAT2 activity by imipramine and mianserin showed no upregulation in VMAT2 mRNA (**Figures 2E-F, 3K-O**). This examination of monoamine transporter gene expression suggests that drug effects of VMAT2 activity upregulation are not through transcriptional regulation.

In stable-transfected HEK293 cells, the total VMAT2 protein levels increased along with an increase in the the mature:immature VMAT2 ratio, suggesting that tricyclics and tetracyclics promote VMAT2 protein maturation (**Figures 3B-G**). This phenomenon of inhibitors promoting transporter maturation has previously been reported for ibogaine, bupropion, and their analogues on DAT, and ibogaine and its analogues on SERT [22, 26–29, 31]. Ibogaine and ibogaine analogues were proposed to act as pharmacological chaperones that bind to the inward-facing conformation of DAT and SERT and stabilize the folding of the transporters. Through stabilization of protein folding, pharmacological chaperones promote maturation of the protein and have an even more substantial effect on folding-deficient disease variants [22, 26, 29]. Since the effects of tricyclics and tetracyclics on VMAT2 upregulation and protein maturation were similar to those of pharmacological chaperones for DAT and SERT, we hypothesize that tricyclics and tetracyclics act as pharmacological chaperones of VMAT2. More investigation is needed to understand the precise mechanism by which tricyclics and tetracyclics promote VMAT2 protein maturation.

The VMAT2 upregulation effect has important clinical implications. It is likely that VMAT2 upregulation is beneficial for mood disorders. Reserpine, an irreversible VMAT1 and VMAT2 inhibitor, was initially used as an anti-hypertensive drug [32]. However, numerous studies have reported depressive mood as an adverse effect of reserpine [14, 15]. The depressive effect of reserpine-induced monoamine depletion was early evidence pointing toward the role of monoamines in depression [4, 33]. Although the current understanding of the pathophysiological mechanisms of mood disorders and depression has changed to include monoamine deficiency, impaired neurogenesis, inflammation, and genetics, VMAT2 inhibition by reserpine remains a valid model of depression in preclinical animal models [34–35]. Reciprocally, mice with genetically elevated VMAT2 level have higher levels of striatal dopamine content and are more resistant to anxiety-like and depressive-like behaviours [36]. Based on human and animal studies, VMAT2 expression levels and activity correlate with depressive mood, and thus upregulation of VMAT2 levels and activity could potentially have antidepressant effects.

The tricyclic and tetracyclics used in our study are well known to have antidepressant effects in humans, however it is unclear to what extent VMAT2 modulation explains their therapeutic effect. An argument against this possibility is the drugs’ mid-micromolar potency for VMAT2 upregulation. However, small molecules that promote VMAT2 maturation would upregulate VMAT2 levels and potentially enhance monoamine transmission. We hypothesize that more potent VMAT2 upregulators could potentially lead to the discovery of a novel class of antidepressants.

Upregulation of VMAT2 could also be beneficial for Parkinson’s disease. A genetic study has found that alleles resulting in VMAT2 overexpression (rs363371 and rs363324) are associated with a lower risk of developing Parkinson’s disease in an Italian population [11]. A study with the Chinese Han population has replicated the result for the rs363371 allele in males, but not the other allele [12]. Another human study has reported gain-of-function VMAT2 haplotypes that result in overexpression of VMAT2 are associated with lower risk of developing Parkinson’s disease in females [13]. Since genetic studies suggest that increased VMAT2 expression is associated with a protective effect for Parkinson’s disease, pharmacological upregulation of VMAT2 levels may also be protective.

Lastly, the rare genetic disease BVMTD, caused by homozygous mutations in the gene coding for VMAT2, could potentially be treated with tricyclics and tetracyclics. In our study, we found that all nine of the missense variants that we tested had VMAT2 functional impairment at baseline. We discovered that imipramine and mianserin upregulated the activity of all variants by 2- to 3-fold compared to vehicle treatment. These preliminary results in cells are very promising as imipramine is FDA-approved for human use, and therefore could be tested for treating BVMTD patients in clinical trials [37]. Most importantly, the two most common recurring variants, P237H and P387L, showed ∼300% upregulation of activity by imipramine compared to their baseline (**Figures 4E, I**). P237H is the most prevalent and most severe BVMTD variant, which has a 53% patient mortality before 13 years of age and currently has no effective treatment options [8]. Upregulation of VMAT2 activity by imipramine could potentially be a promising treatment for these BVMTD patients.

In conclusion, our study is the first to report that clinically-relevant tricyclic and tetracyclic antidepressants inhibit VMAT2 activity with acute treatment but upregulate VMAT2 activity with sustained treatment by promoting VMAT2 protein maturation. The upregulation effect was greater in the endogenous BE(2)-M17 cell system than in the HEK expression system. However, HEK cells were useful to detect the drugs’ effects on protein maturation using western blot to identify glycosylated protein forms. Imipramine and mianserin, representative tricyclic and tetracyclics, rescued fatal disease-causing variants of VMAT2.

A limitation of our study is that our findings are restricted to cell systems, therefore further investigation *in vivo* is necessary to ascertain whether tricyclics or tetracyclics also upregulate VMAT2 *in vivo*. Tricyclic and tetracyclic antidepressants have long been used in the clinic for the treatment of depression in humans, thus studies involving positron emission tomography (PET) imaging could be conducted to investigate if the VMAT2 protein upregulation phenomenon could be observed in patients who are taking tricyclic or tetracyclic antidepressants. In addition, our study identified imipramine and mianserin as two promising therapeutic candidates for BVMTD patients, warranting clinical trials for this fatal disease.

## Acknowledgments

We would like to acknowledge Dr. Aurora Martinez for helpful scientific discussion on the manuscript and the Flow Cytometry Facility at the University of Toronto for helping us acquire flow cytometry data.

## Author Contributions

XW, IM, PSBF, SIS, MLB, TVL, AJR, GWM and AS designed the study; XW, IM, PSBF, SIS, MLB, TVL, and AJR conducted experiments to acquire data; XW wrote the manuscript draft; all other authors revised, finalized and approved the manuscript.

