## Supplementary materials for "Tricyclic and tetracyclic antidepressants upregulate VMAT2 activity and rescue disease-causing VMAT2 variants"

### Supplementary Tables

Table S1. Reagent catalogue and supplier information

Table S2. Site-directed mutagenesis PCR primers used for constructing VMAT2 BVMTD variants.

Table S3. RT-qPCR primers used for quantitative assessment of gene expression.

### Supplementary Figures

Figure S1. Example gating for BE(2)-M17 cell FFN206 uptake flow cytometry

Figure S2. Epifluorescent microscopy of FFN206 colocalization with VMAT2-mCherry.

Figure S3. Dose-response of 30-minute incubation with TBZ on VMAT2 activity.

Figure S4. Western blot analysis of VMAT2 deglycosylation.

Figure S5. Western blot analysis of 18-hour sustained imipramine or mianserin treatment on VMAT2 protein levels.

Figure S6. Western blot analysis of 18-hour sustained imipramine or mianserin treatment on YFP-DAT protein levels.

**Table S1. Reagent catalogue and supplier information**

| Categories | Reagent (catalogue) | Supplier |
|---|---|---|
| Cell lines | HEK293 (CRL-1573) | ATCC |
| | HEK293T (CRL-3216) | ATCC |
| | BE(2)-M17 (CRL-2267) | ATCC |
| Reagents | Penicillin and streptomycin solution, 100X (PST999) | Bioshop |
| | G418 sulfate powder (GEN418.5) | Bioshop |
| | CELLSTAR® 96 well Microplates, black plate with black bottom (655086) | Greiner Bio-One |
| | Dulbecco′s Phosphate Buffered Saline, with MgCl2 and CaCl2 (D8662) | Sigma-Aldrich |
| | 6-well plate, clear (3516) | Corning |
| | Trypsin-EDTA (0.25%) (25200056) | Gibco |
| | Pierce™ BCA Protein Assay Kits (23225) | ThermoFisher Scientific |
| | Endo H (P0702S) | New England Biolabs, Inc. |
| | PNGase F (P0704S) | New England Biolabs, Inc. |
| | NuPAGE™ 4 to 12% Bis-Tris Mini Protein Gel (NP0322BOX) | Invitrogen |
| | Revert™ 700 Total Protein Stain | LI-COR, Inc. |
| | rabbit-anti-GFP antibody (A11122) | Invitrogen |
| | IRDye® 800CW Goat anti-Rabbit IgG (926-32211) | LI-COR, Inc. |
| | Lipofectamine™ 3000 | Invitrogen |
| | TRIzol Reagent | Invitrogen |
| | SuperScript™ IV VILO™ Master Mix with ezDNase™ Enzyme | Invitrogen |
| | PowerUp™ SYBR™ Green Master Mix | Applied Biosystems |
| Chemicals | FFN206 dihydrochloride (5043) | Tocris Bioscience |
| | Tetrabenazine (T284000) | Toronto Research Chemicals |
| | Reserpine (R144600) | Toronto Research Chemicals |
| | Mianserin hydrochloride (M341500) | Toronto Research Chemicals |
| | Amitriptyline hydrochloride (A633350) | Toronto Research Chemicals |

| | | |
|---|---|---|
| | Amoxapine (A634230) | Toronto Research Chemicals |
| | Desipramine hydrochloride (D290050) | Toronto Research Chemicals |
| | Doxepin hydrochloride (D550000) | Toronto Research Chemicals |
| | Imipramine hydrochloride (I465980) | Toronto Research Chemicals |
| | Nortriptyline hydrochloride (N837000) | Toronto Research Chemicals |
| | Trimipramine maleate salt (T799000) | Toronto Research Chemicals |
| | Protriptyline hydrochloride (P838875) | Toronto Research Chemicals |

**Table S2. Site-directed mutagenesis PCR primers used for constructing VMAT2 BVMTD variants.**

| Construct | Mutation | Forward primer (5' to 3') | Reverse primer (5' to 3') |
|---|---|---|---|
| pcDNA3.1-hVMAT2-P42L | P42L | gctcactgtcgtggtcctcatcatcccaagttatc | gataacttgggatgatgaggaccacgacagtgagc |
| pcDNA3.1-hVMAT2-I43F | I43F | cactgtcgtggtccccttcatcccaagttatct | agataacttgggatgaaggggaccacgacagtg |
| pcDNA3.1-hVMAT2-P237H | P237H | ttagtgggcccccacttcgggagtgtg | cacactcccgaagtgggggcccactaa |
| pcDNA3.1-hVMAT2-A309V | A309V | caaacatgggcatcgtcatgctggagccagc | gctggctccagcatgacgatgcccatgtttg |
| pcDNA3.1-hVMAT2-P316A | P316A | ggagccagccctggccatctggatgat | atcatccagatggccagggctggctcc |
| pcDNA3.1-hVMAT2-A333D | A333D | gcagctgggcgttgacttcttgccagcta | tagctggcaagaagtcaacgcccagctgc |
| pcDNA3.1-hVMAT2-P387L | P387L | catttatggactcatagctctgaactttggagttggttttgc | gcaaaaccaactccaaagttcggagctatgagtccataaatg |
| pcDNA3.1-hVMAT2-Y418C | Y418C | gcggcacgtgtccgtctgtgggagtgt | acactcccacagacggacacgtgccgc |
| pcDNA3.1-hVMAT2-G436S | G436S | tttgtatggggtatgctataagtccttctgctggtg | caccagcagaaggacttatagcataccccatacaaa |

**Table S3. RT-qPCR primers used for quantitative assessment of gene expression.**

| Gene | Forward (5' to 3') | Reverse (5' to 3') | Sequence |
|---|---|---|---|
| hGAPDH | GTCTCCTCTGACTTCAACAGCG | ACCACCCTGTTGCTGTAGCCAA | NM_001256799.3 |
| mVMAT2 | CCTCTTACGACCTTGCTGAAGG | GCTGCCACTTTCGGGAACACAT | BC078449.1 |
| hVMAT2 | GCTATGCCTTCCTGCTGATTGC | CCAAGGCGATTCCCATGACGTT | NM_003054.6 |
| hVMAT1 | AGGTTTCTTGGAGGAAGAGATTAC | ATCCAATCCTGTTGGTGAGAG | NM_003053.4 |
| hDAT | CCTCAACGACACTTTTGGGACC | AGTAGAGCAGCACGATGACCAG | NM_001044.5 |
| hSERT | TCACAGTGCTCGGTTACATGGC | GAAAGTGGACGCTGGCATGTTG | NM_001045.6 |
| hNET | CAGGTTCAGCAACGACATCCAG | GTCGTAGGTGAGTGGCTTGAAG | NM_001043.3 |

**GAPDH:** Glyceraldehyde 3-phosphate dehydrogenase; **VMAT:** Vesicular Monoamine Transporter; **DAT:** dopamine transporter; **SERT:** Serotonin transporter; **NET:** Norepinephrine transporter.

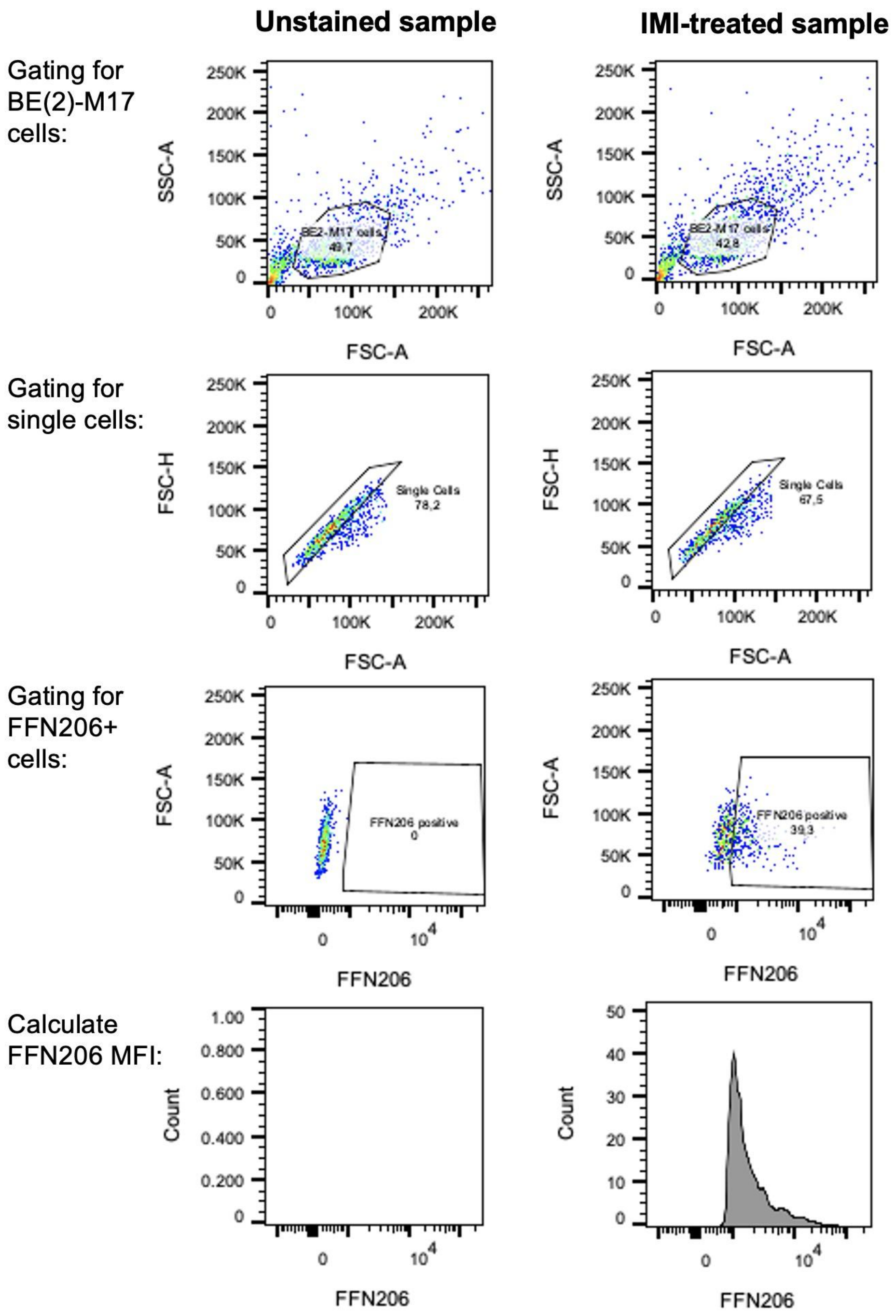


**Figure S1. Example gating for BE(2)-M17 cell FFN206 uptake flow cytometry.** Gatings: BE(2)-M17 cells/single cells/FFN206 positive cells. Mean fluorescence intensity (MFI) was quantified for FFN206 positive cells. Left: unstained sample (no FFN206); right: imipramine-treated sample with FFN206.

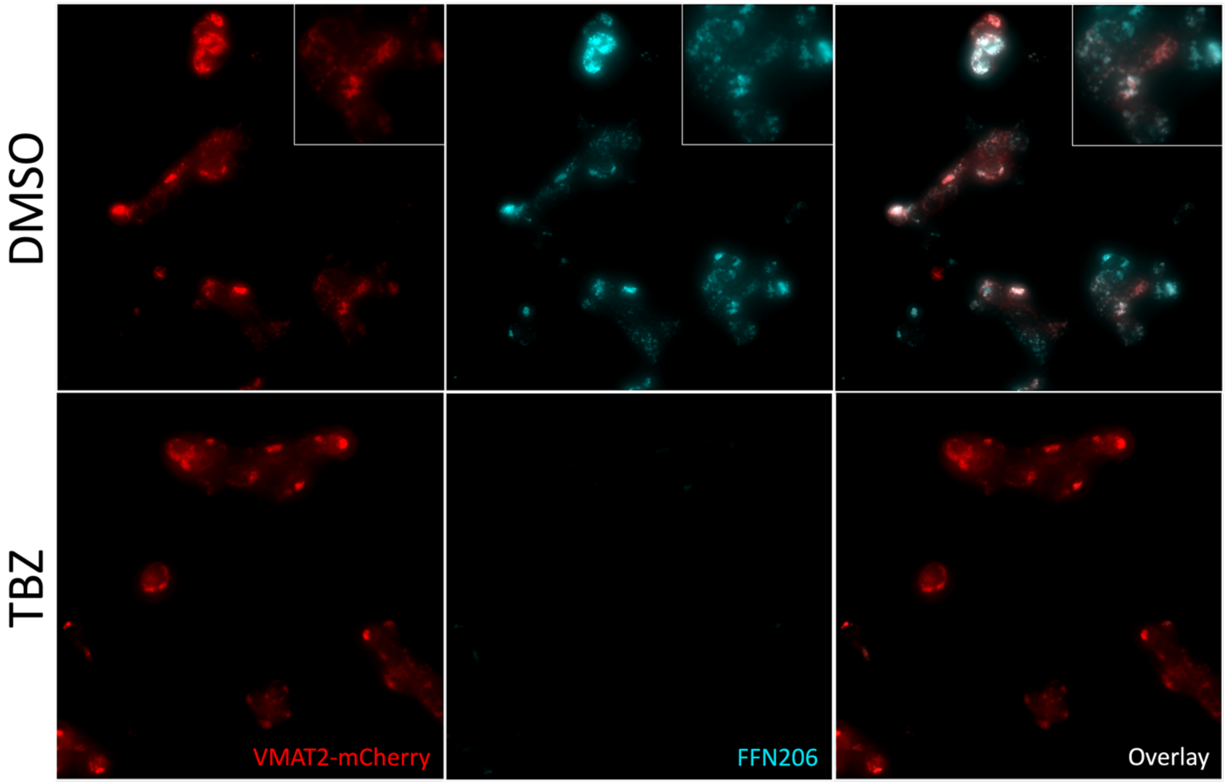


**Figure S2. Epifluorescent microscopy of VMAT2-mCherry and FFN206.** Red: VMAT2-mCherry; Cyan: FFN206; White: Overlay of VMAT2-mCherry and FFN206. VMAT2 and FFN206 colocalize in intracellular compartments.

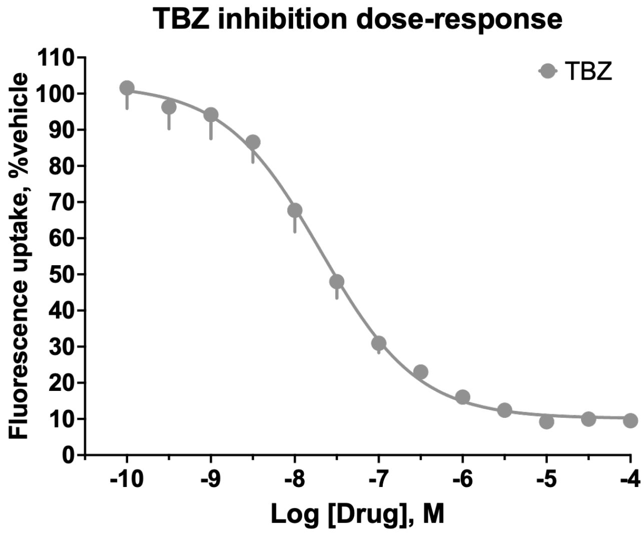


**Figure S3. Dose-response of 30-minute incubation with TBZ on VMAT2 activity in HEK-VMAT2 cells (n = 10).**

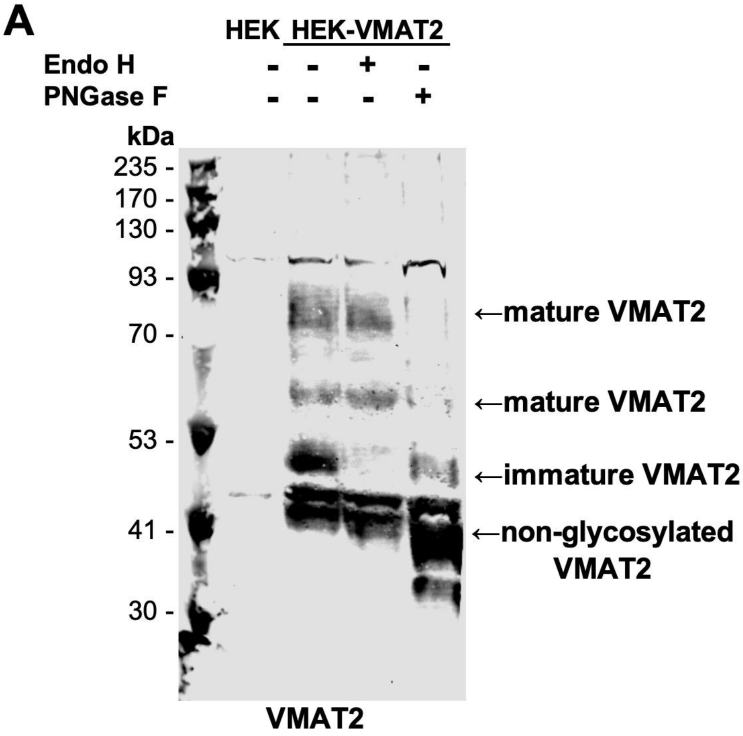


**Figure S4. Western blot analysis of VMAT2 deglycosylation. (A)** Western blot of endoglycosidase H (EndoH) and peptide:N-glycosidase F (PNGase F) digested VMAT2 protein revealing different trafficking stages of the VMAT2 protein. **(B)** Total protein stain of blot A as a loading control.

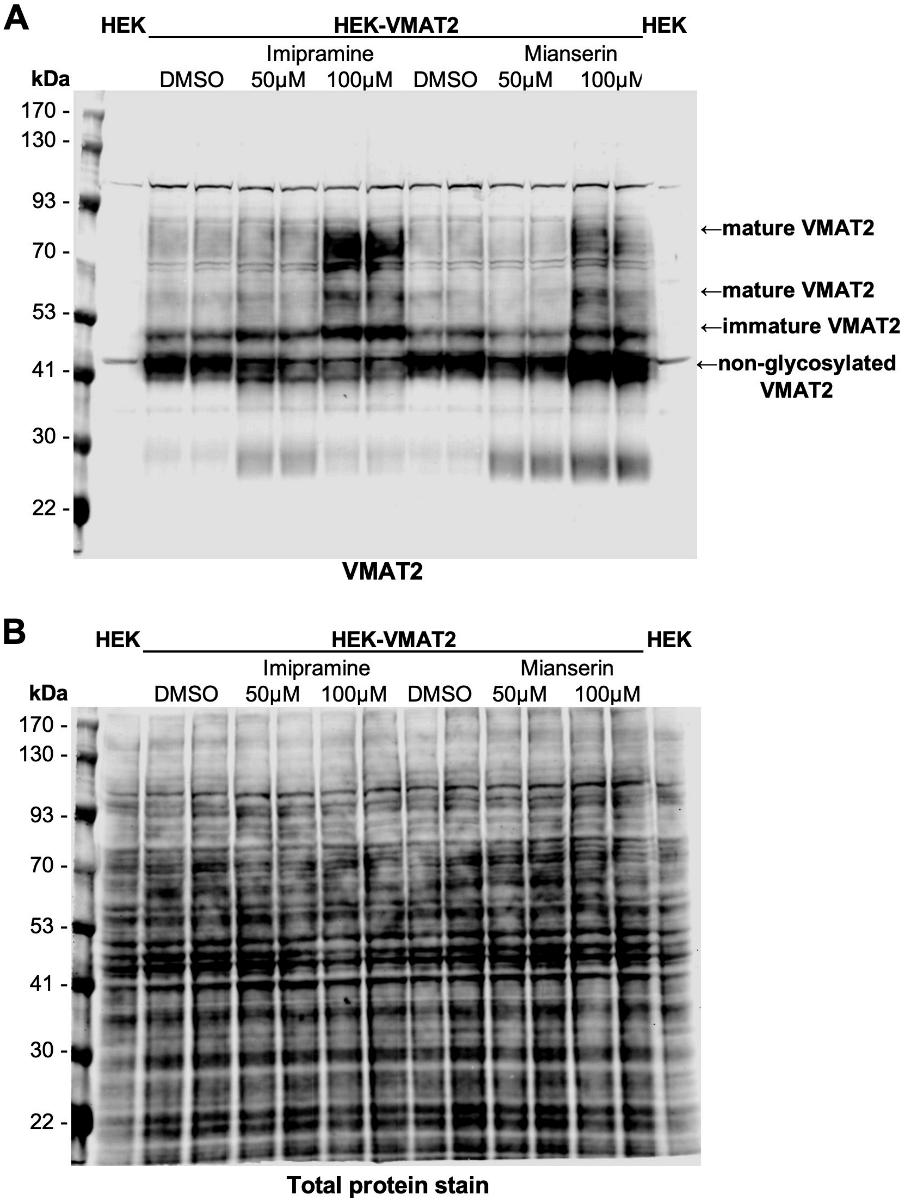


**Figure S5. Western blot analysis of 18-hour sustained imipramine or mianserin treatment on VMAT2 protein levels. (A)** Representative western blot of 18-hour imipramine or mianserin incubation on VMAT2 protein in HEK-VMAT2 cells. Duplicate wells (n=1) displayed. **(B)** Total protein stain of blot A as a loading control.

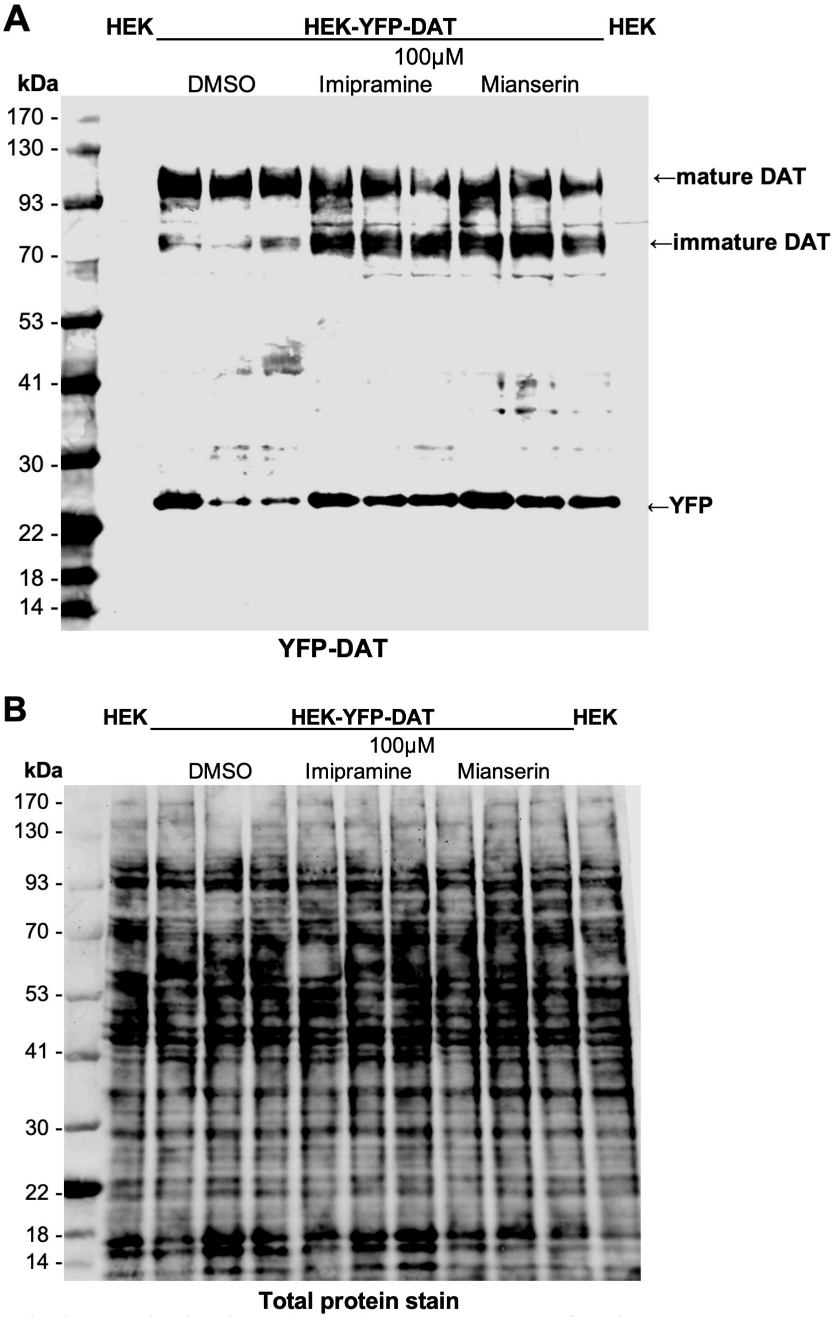


**Figure S6. Western blot analysis of 18-hour sustained imipramine or mianserin treatment on YFP-DAT protein levels. (A)** Western blot of 18-hour 100 µM imipramine and mianserin incubation on DAT protein in HEK-YFP-DAT cells (n = 3, each well represents an independent experiment). **(B)** Total protein stain of blot A as a loading control.
